# Temporal Progression of Prior Precision, Surprisal, Model Updating and Tone Adaptation in Human Auditory Cortices

**DOI:** 10.64898/2026.10.25.761658

**Authors:** Jorie van Haren, Floris P. de Lange, Sonja A. Kotz, Federico De Martino

## Abstract

Auditory perception emerges from the continuous integration of incoming acoustic signals with statistical predictions, a process complemented by simpler local adaptation. Yet, how all these concurrent mechanisms unfold in time remains underexplored. Using magnetoencephalography (MEG) in participants listening to stochastic tone sequences with periodically shifting inferential statistics, we modelled the temporal, spatial, and spectral dynamics of stimulus-specific repetition suppression, surprisal, model updating, and prior precision. By concurrently tracking these computational parameters within this dynamic auditory environment, we revealed a distinct chronological hierarchy governing predictive and adaptive processing. Prior precision states - reflecting expectations - emerged earliest, followed by rapid and concurrent repetition suppression and surprisal - reflecting local acoustic attenuation and global mismatch detection. These initial dynamics culminated in subsequent, distributed model updates within fronto-parietal networks. Spectrally, these components mapped onto diverging oscillatory fingerprints, ranging from theta-alpha modulation during adaptation to beta suppression during surprisal and broad spectral dampening during precision weighting. Together, these findings demonstrate that the brain navigates uncertain auditory environments through a cascade of neural processes, where fast, local sensory gating and mismatch signals precede the hierarchical revision of internal models.

## 1 Introduction

Auditory perception is shaped by the brain’s continuous integration of incoming acoustic signals with contextual cues spanning multiple timescales [1, 2]. These contextual cues are used to forecast future events, allowing us to gauge whether our observations match our internal predictions. In turn, novel information - representing a departure from these same predictions - is used to update our predictive models, resulting in a continuously evolving inference of the auditory environment [3, 4]. These forward-looking processes do not act in isolation; they are aided by local adaptation mechanisms that attenuate neural responses to recent acoustic history, resulting in observed repetition suppression [5, 6, 7, 8]. Because our predictions often coincide with repetitions of low-level features - such as the acoustic frequency of a sound - this attenuation can strongly mimic prediction-related modulation at the neural level [7, 9, 10, 11].

While prior work has hinted at when predictive processes occur - such as demonstrating that surprisal precedes model updating [12], or that acoustic deviations elicit a mismatch negativity (MMN) response [13] - early research often investigated these components in isolation. To address the limitations of isolated designs, subsequent studies have sought to decouple interacting variables, utilising orthogonal factorial designs to separate repetition from expectation [14, 9], or employing computational models to distinguish instantaneous surprisal from continuous model updating [15]. However, because these approaches frequently treat expectations as discrete categorical states or model only a subset of variables, they struggle to fully capture the continuous dynamics present in statistical learning. This partial modelling makes it difficult to map specific neural signatures to their correct cognitive processes. Without testing related variables concurrently, measured components risk absorbing the neural variance of omitted processes, leading to statistical confounding and misattribution [16]. Consequently, how local adaptation, parametric surprisal, model updating, and prior precision operate in tandem within a continuously evolving statistical environment remains underexplored [7].

Theoretical frameworks and neurophysiological evidence suggest a specific temporal hierarchy governing these predictive and adaptive mechanisms. Because the precision of our prior expectations - reflecting the confidence in our internal model - dictates the weighting of mismatches derived from incoming sensory evidence, this top-down state is expected to emerge earliest [17]. This is likely followed by repetition suppression, a rapid mechanism that integrates with initial sensory engagement within the feedforward pathway [18, 11]. Subsequently, the initial encoding of surprisal serves as a global signal reflecting the overall magnitude of unexpectedness [19]. In line with dual-opponent accounts of predictive processing, where an initial bias toward expected structure transitions to prioritise informative deviations [20], this early surprisal signal likely precedes the resulting model updates [12]. However, because the evidence supporting this chronological cascade relies on experiments that attempt to isolate individual components (e.g. mismatches or predictions), the stability of this exact sequential hierarchy within a continuous, stochastically evolving environment remains uncertain.

Similarly, when considering these components in tandem, the specific spectral bands they occupy remain unclear. Current predictive coding frameworks propose a spectral dissociation between computations such as predictions and model updating, though these mappings remain debated and often vary with the experimental paradigm. Generally, top-down predictions and their associated prior precision are linked to lower-frequency alpha and beta oscillations [21, 17], which are thought to reflect the maintenance and propagation of the internal model. Conversely, bottom-up prediction errors and surprisal are typically linked to higher-frequency gamma-band activity [21, 17, 22], alongside theta oscillations commonly observed during the MMN response [23, 24]. However, when these signals are investigated in their role in driving active model updating, their spectral signatures have also been found to extend into the alpha and beta bands, with alpha related to the bidirectional interaction of hierarchical cortical levels [25] and attenuated beta indicating a departure from the maintained model [26]. Meanwhile, repetition suppression leading to adaptation often manifests as attenuated gamma power [7], yet concurrent modulations in theta oscillations point to a more complex spectral signature [27]. Given the established difficulty of studying these mechanisms in isolation and the risk of conflating overlapping spectral signatures, their precise frequency allocation remains unresolved, further emphasising the need to model these processes concurrently.

In the current study, we investigate the temporal order and latency of computationally modelled parametric estimates of predictive and adaptive mechanisms within a statistically evolving environment. Specifically, participants passively listened to sequences of pure tones sampled stochastically from Gaussian distributions while their ongoing neuromagnetic responses were recorded using magnetoencephalography (MEG). The sampling probabilities of the Gaussian distributions changed over time, forcing listeners to update their internal model continuously. We then relied on time-shifted regressions to see how normatively modelled - graded - estimates of repetition suppression, surprisal, model updating, and prior precision evolved temporally. The same procedure was applied within the spectrotemporal representation of the recorded signals to investigate the spectrotemporal power dynamics. Our results reveal a distinct chronological hierarchy governing how the brain navigates probabilistic acoustic environments: prior precision states, reflecting expectations, emerge earliest, followed by rapid, concurrent stimulus-specific adaptation and mismatch detection, and culminating in delayed model updates.

## 2 Results

Thirty-two participants passively listened to stochastic tone sequences while magnetoencephalography (MEG) was recorded. Sequences comprised pure tones sampled from two Gaussian distributions with shifting probabilities, designed to maximise the decorrelation between stimulus-specific adaptation and surprisal (*r ≈* 0.68, compared to *r ≈* 0.99 in classical oddball paradigms; see Figure 1-Top). We investigated the temporal dynamics of repetition suppression, surprisal, model updating, and prior precision. Repetition suppression was modelled as a frequency-specific attenuation governed by a double-exponential decay, which captures both the rapid and sustained adaptation to similar tones over recent trials ([8]; see Methods 4.5.1). Concurrently, predictive processes were estimated using the D-REX model, which generates probabilistic Bayesian forecasts by inferring the statistical regularities of the sound sequence, thereby deriving both stimulus surprisal (i.e., the *−* log(*p*)) and prior precision (i.e., the certainty of the prior predictions; [28]; see Methods 4.5.2). Building on this, model updating (i.e., the extent of belief revision) was quantified continuously as the integrated difference between the model’s prior expectations and the updated posterior distributions following each tone ([29]; see Methods 4.5.3). Crucially, evaluating the multiple collinearity of this full model space confirmed that at least 49% of the variance for every computational parameter remained unique and orthogonal to the rest of the model (for all pairwise correlations and unique variance metrics, see Supplementary Table **S1**).

**Figure 1:**
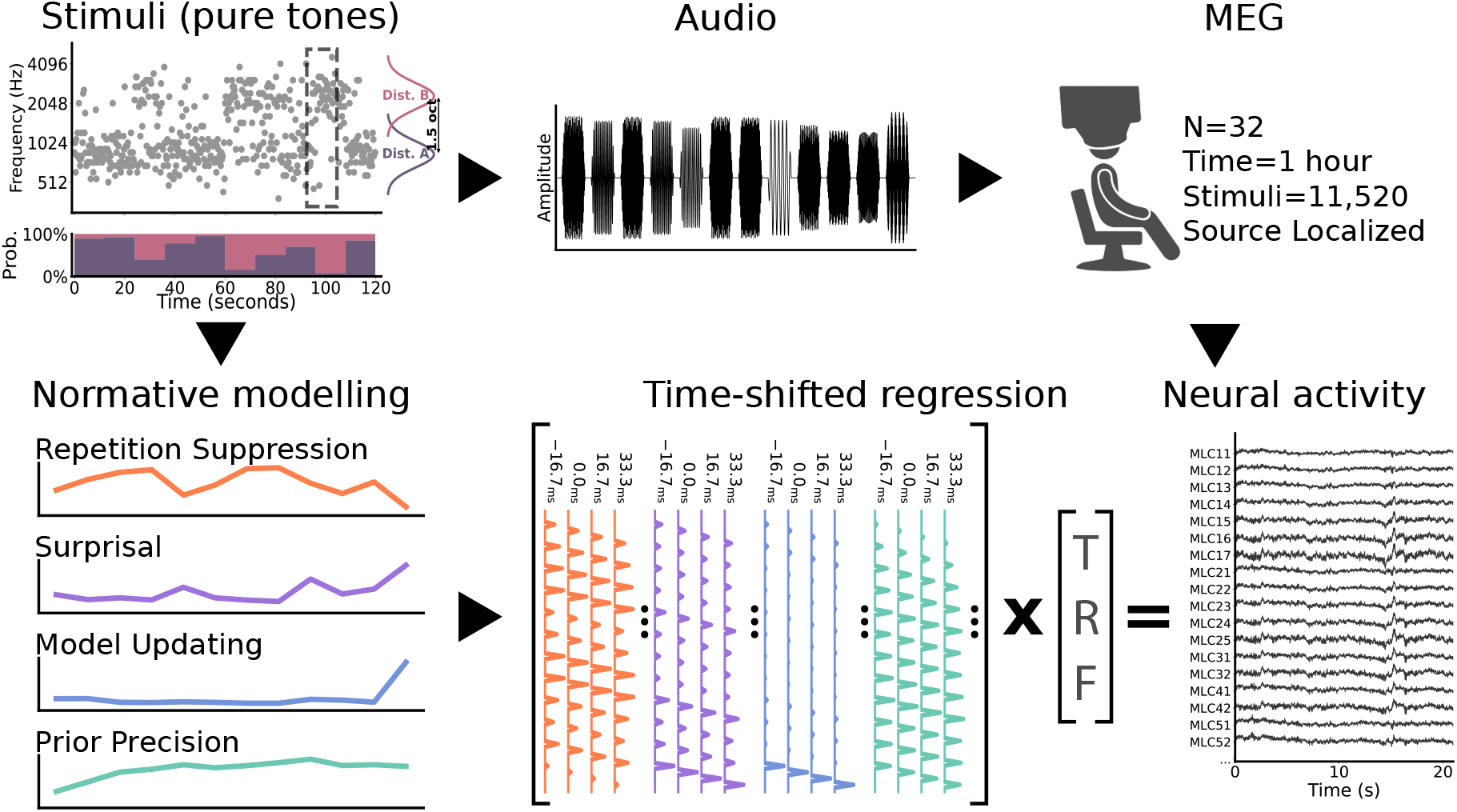
Schematic of the experimental and analytical framework. (Top) Participants passively listened to stochastic tone sequences, while their brain activity was measured using MEG. (Bottom) The tone sequences participants listened to were used in normative modelling approaches to estimate the expected amounts of repetition suppression, surprisal, model updating, and prior precision for each tone. Repetition suppression was modelled using a frequency-specific double-exponential decay, surprisal as the unexpectedness of a tone given its probabilistic Bayesian forecast (Shannon; *−* log(*p*)), model updating as the integrated difference between prior expectations and updated posteriors, and prior precision as the inferred certainty of the evolving statistical expectations. We estimate the temporal response functions for the different features using time-shifted (FIR) linear regressions on the MEG data, while controlling for low-level acoustic factors.

To account for the rapid, continuous presentation of the 11,520 stimuli in total and the resulting overlap of neural responses within our temporal window of interest (*−* 50 ms to 567 ms; chosen to be coprime to the 250 ms stimulus interval to naturally jitter the overlap of successive trial responses, which helps disentangle their temporal dynamics; see Methods 4.6.2), we employed a linear deconvolution framework based on time-shifted finite impulse response (FIR) models (see Figure 1-Bottom). The regression model also included nuisance regressors for basic tone-onset effects and allowed us to estimate the temporal dynamics of each predictor of interest (repetition suppression, surprisal, model updating, and prior precision) from the associated beta coefficients. The resulting Temporal Response Functions (TRFs) were subsequently source-localised using individually constructed, anatomically motivated common filters (LCMV).

### 2.1 Neural Responses are Modulated by Continuous Predictive and Adaptive Processes

We first tested for general evidence of concurrent predictive and adaptive processing. We reasoned that if the brain continuously tracks statistical regularities, neural responses to tones should be sensitive to violations of predictions (surprisal), the updating of internal models (model updating), and changes in the uncertainty of these models (prior precision), while simultaneously being modulated by local acoustic attenuation (repetition suppression).

To investigate where and when the response functions captured significant neural modulation, we performed mass univariate tests on the source-localised TRF coefficients (i.e., beta weights), testing them against a non-parametric null distribution obtained by block-permuting and phase-decoupling the target regressors (see Methods 4.6.2). Multiple-comparison correction across cortical areas and time (from -50 ms to 567 ms) was achieved using threshold-free cluster enhancement (TFCE), with statistical significance assessed at a conservative *α <* 0.01 to further control for the evaluation of four distinct parameters.

This analysis revealed significant spatiotemporal clusters for all regressors, confirming that neural activity was independently modulated by each computational parameter (all *p <* 0.01; TFCE-corrected). To summarise the temporal extent of the effects, we extracted temporal peaks by evaluating the integrated statistical strength (i.e. area under the curve; AUC) of the clusters across a 100-ms rolling window (see Methods 4.9.1). Both predictive and adaptive computations elicited sustained responses spanning most of the temporal window. For repetition suppression, the significant modulation was driven by a cluster spanning the full temporal window from -50 ms to 567 ms, a continuous profile consistent with the ongoing attenuation of prior stimulus history (see Figure 2a). This effect peaked in statistical power at 183 ms, an early latency characteristic of neural gain attenuation. Similarly, the significant effect of surprisal was most compatible with a broad continuous cluster spanning from 50 ms to 567 ms and peaking earlier at 117 ms (see Figure 2b), a profile consistent with its stimulus-dependent nature. The significant effect of model updating was driven by an effect starting at 33 ms and continuing until 567 ms, but peaking later at 200 ms (see Figure 2c). Finally, prior precision evoked a distinct temporal profile, with the significant difference driven by an effect also spanning the full epoch from -50 ms to 567 ms but reaching its peak at 167 ms (see Figure 2d), a latency consistent with its reliance on established prior expectations. Together, these findings provide clear evidence for concurrent predictive and adaptive processing, indicating that continuous neural responses are modulated by both local acoustic history and hierarchical statistical predictions.

**Figure 2:**
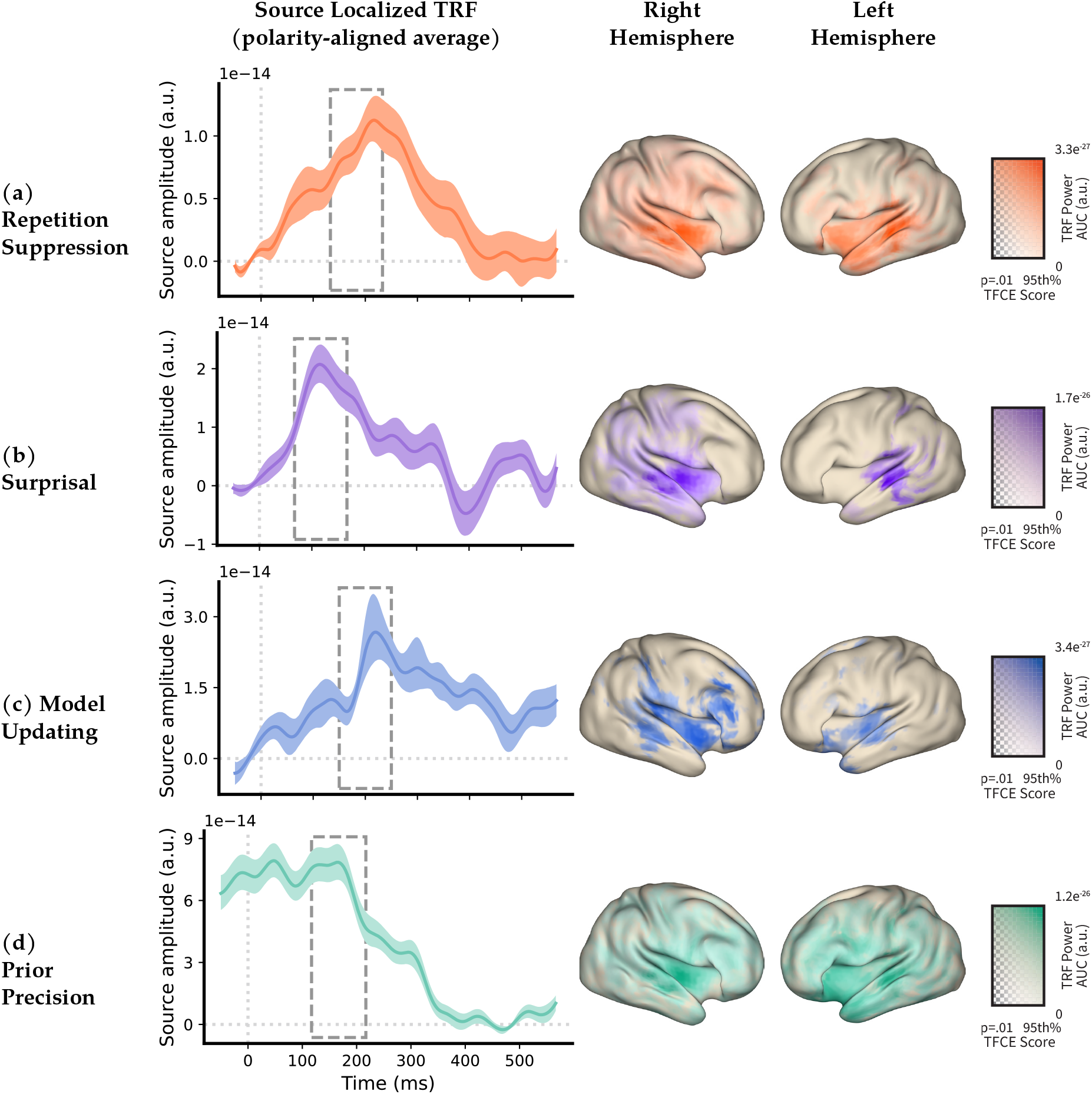
Spatiotemporal signatures of predictive and adaptive processing components. (a) Repetition suppression, (b) surprisal, (c) model updating, and (d) prior precision. Left: Polarity-aligned, source-localised temporal response functions (TRFs) derived from time-shifted finite impulse response (FIR) regressions. Right: TRF waveforms represent the signal extracted from the spatial region of interest containing the greatest statistical strength for each respective component (left middle temporal gyrus for repetition suppression, surprisal, and model updating; right middle frontal gyrus for prior precision) across participants. Shaded areas indicate bootstrapped 95% confidence intervals. Dashed boxes denote a 100-ms integration window centred on the temporal peak of the underlying significant spatiotemporal Threshold-Free Cluster Enhancement (TFCE) cluster. Note that this window reflects the period of maximal statistical significance across the spatiotemporal cluster, rather than the peak amplitude of the plotted group-average waveform. Colour hue represents the maximum squared coefficient (power) extracted from the respective 100-ms peak statistical window. Transparency (alpha) is mapped to the underlying TFCE statistics, scaled between the cluster-defining significance threshold (*p* = 0.01, fully transparent) and the 95th percentile of significant TFCE values (fully opaque).

### 2.2 Spatial signatures of temporal repetition suppression and surprisal versus downstream model updating and widespread prior precision networks

Having established that predictive context and tone repetition modulate overall neural responses, we characterised these effects in space. To identify the primary anatomical basis of each effect, we parcellated the source-localised clusters identified above using the AAL atlas, determining the dominant regions of interest (ROIs) based on their statistical strength - defined as the cumulative TFCE statistics across the full temporal cluster (see Methods 4.9.1). Alongside identifying these core anatomical regions, we quantified the broader spatial spread of each effect by reporting its ‘footprint at peak’ - defined as the percentage of the intracranial source space significantly active within a 100-ms window centred on the cluster’s temporal peak (for an overview of all footprints and anatomical dispersion metrics, see Supplementary Table **S3**).

For repetition suppression (temporal window: -50 – 567 ms; footprint at peak: *∼* 78% of the intracranial source space), TRF power was broadly clustered within and around auditory regions, with the effect being concentrated on the superior (right *p*_*min*_ *<* 0.001, left *p*_*min*_ *<* 0.001) and middle temporal gyri (right *p*_*min*_ *<* 0.001, left *p*_*min*_ *<* 0.001; see Figure 3a). These bilateral temporal effects reflect the engagement of established auditory processing pathways, yielding a spatial distribution consistent with local adaptation to recent acoustic history.

**Figure 3:**
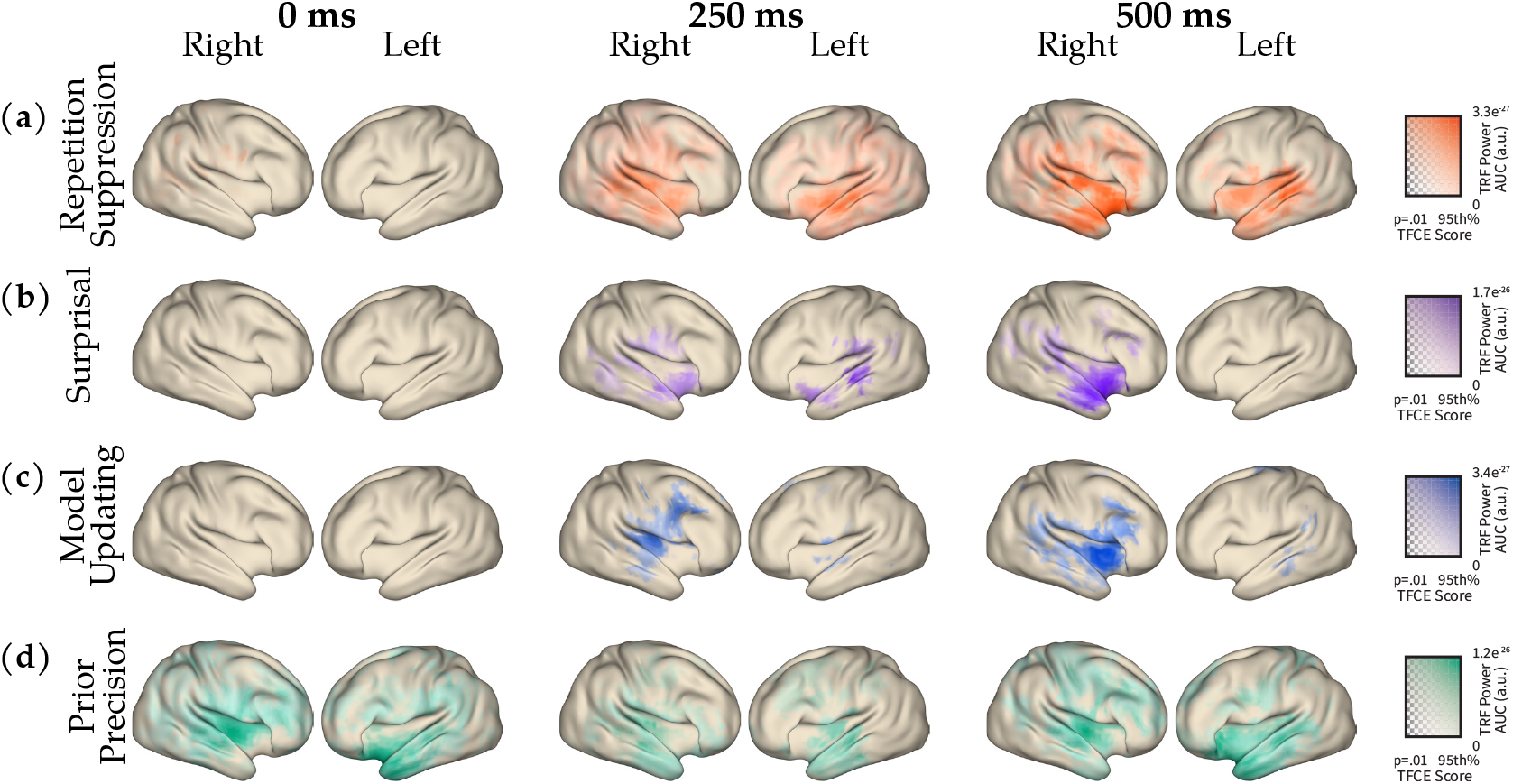
Spatiotemporal evolution of cortical power responses. Cortical surface maps display the distribution of source-localised MEG power for four experimental components: (a) Repetition suppression, (b) surprisal, (c) model updating, and (d) prior precision. Columns represent 50ms time windows centred at 0 (-25 - 25ms), 250 (225 - 275ms), and 500 ms (475 - 525ms) relative to stimulus onset, for both right and left hemispheres. Colour maps reflect the integrated statistical strength (area under the curve; AUC) of the TRF power within each time window. Colour-bar hue indicates the magnitude ranges specific to each component’s statistical map (TRF Power AUC), while alpha reflects the statistical significance (TFCE Score), matched across all time points within that condition.

Surprisal (temporal window: 50 – 567 ms; footprint at peak: *∼* 31% of the intracranial source space) was similarly centred around primary and associative auditory cortices, with clustered statistical strength centred on the superior (right *p*_*min*_ *<* 0.001, left *p*_*min*_ = 0.001) and middle temporal gyri (right *p*_*min*_ *<* 0.001, left *p*_*min*_ = 0.001; see Figure 3b). In contrast to the more diffuse spread of repetition suppression, the effects of surprisal were more focal within these core auditory regions, additionally engaging earlier auditory areas such as Heschl’s gyrus alongside secondary clustering observed around the temporal poles.

In contrast to surprisal, model updating signals exhibited a more distributed and temporally hierarchical organisation. While model updating partially shared a spatio-temporal locus with the other parameters, its initial cortical representation was less reliant on primary sensory regions. Whereas surprisal maximally recruited primary auditory areas like Heschl’s gyrus, the early temporal window of model updating (50 – 267 ms; footprint at peak: *∼* 38% of the intracranial source space) was more prominent in associative auditory and insular regions (right *p*_*min*_ *<* 0.001, left *p*_*min*_ = 0.001). Following this early temporal response, model updating recruited a secondary cortical network (367 - 567 ms) that extended significantly into the lateral prefrontal cortex (right *p*_*min*_ *<* 0.001, left *p*_*min*_ = 0.002; see Figure 3c), as well as broader parietal regions, particularly during the later processing stages (*>* 350 ms). This transition - from an early associative temporal locus to a later, distributed fronto-parietal signature - suggests that model updating moves beyond the initial sensory mismatch detection characteristic of surprisal, progressing into integrated, higher-order evaluative processing within frontal areas.

Finally, prior precision displayed a distinct topology. Rather than being confined to the core temporal network like repetition suppression or surprisal, the spatial signature of prior precision was highly distributed. Specifically, prior precision (temporal window: -50 – 567 ms; footprint at peak: *∼* 92% of the intracranial source space) recruited a widespread network encompassing both associative temporal regions and extensive portions of the prefrontal cortex (right *p*_*min*_ *<* 0.001, left *p*_*min*_ *<* 0.001; see Figure 3d). Moreover, prior precision uniquely recruited an additional, spatially distinct locus emerging later in the epoch (533 - 567 ms) localised to the cerebellum (left *p*_*min*_ = 0.009). This spatial dissociation suggests that while distributed fronto-cortical networks handle the primary precision-weighting signals, the cerebellum contributes uniquely to a possible later model-updating signal.

### 2.3 Surprisal Precedes Model Updating in Predictive Processing

While the preceding spatial analyses clarified how repetition suppression, surprisal, model updating, and prior precision differentially recruit temporal and fronto-parietal networks, we next investigated how these effects unfold over time. Because our deconvolution framework estimates these parameters concurrently, we can directly compare their latencies to determine whether the brain computes these distinct features simultaneously or sequentially. This chronological mapping allows us to test a core assumption of hierarchical predictive processing: that rapid, lower-level mismatch detection strictly precedes higher-order model updating.

Investigating whether these predictions unfold over different timescales required evaluating the relative timings of the components within the dominant clusters identified above (i.e., the specific clusters exhibiting the highest TFCE statistical strength; for comparison of the secondary and lateralised clusters see Supplement **S2**). To ensure our latency estimates were physiologically constrained and robust to baseline slow-wave shifts, we restricted our peak search exclusively to the significant temporal windows defining these clusters. By defining component latencies using a 50% relative amplitude criterion (half-maximum) computed relative to the signal onset within that cluster window, we isolated the transient onset of the neural response driven by each specific computational process (i.e., repetition suppression, surprisal, model updating, and prior precision). These latency estimates were compared using jackknife-based latency *t*-tests ([30]; for an overview of all latencies, see Supplementary Table **S2**; see Methods 4.11).

This analysis revealed a temporal separation between rapid, lower-level mismatch detection and delayed, higher-order hierarchical updates. Repetition suppression and surprisal evoked rapid, temporally aligned responses, reaching their relative half-maximums at 74.1 ms and 80.2 ms respectively, with no statistical difference in their timing (*t*_31_ = *−* 0.29, *p*_FDR_ = .773). In contrast, the temporal signatures of higher-order updates occurred substantially later. Both surprisal (*t*_31_ = *−* 20.95, *p*_FDR_ *<* .001) and repetition suppression (*t*_31_ = *−* 5.56, *p*_FDR_ *<* .001) reached half-maximum significantly earlier than model updating (*M* = 188.8 ms). Prior precision represented a pre-existing state, reaching its relative half-maximum significantly earlier than all other components just before stimulus onset (*−* 12.8 ms): prior to surprisal (*t*_31_ = *−* 6.11, *p*_FDR_ *<* .001), repetition suppression (*t*_31_ = *−* 3.49, *p*_FDR_ = .002), and model updating (*t*_31_ = *−* 14.12, *p*_FDR_ *<* .001). However, because prior precision exhibits a sustained baseline shift starting before stimulus onset, evaluating its later offset latency (178.7 ms) - which captures the decay of this prior-dependent state - may provide a more informative metric for comparing the completion of these processing stages.

The decay of prior precision (*M* = 178.7 ms) significantly preceded the offset tail of repetition suppression (*M* = 311.0 ms; *t*_31_ = *−* 8.31, *p*_FDR_ *<* .001) and model updating (*M* = 398.3 ms; *t*_31_ = *−* 2.72, *p*_FDR_ = .021). However, it did not significantly precede the offset tail of surprisal (*M* = 192.4 ms; *t*_31_ = *−* 1.52, *p*_FDR_ = .166). Similarly, the surprisal effect resolved significantly earlier than both repetition suppression (*t*_31_ = *−* 7.88, *p*_FDR_ *<* .001) and model updating (*t*_31_ = *−* 2.52, *p*_FDR_ = .026), which showed no statistical difference in the timing of their extended processing tails (*t*_31_ = *−* 1.09, *p*_FDR_ = .286). Together, this indicates that while the decay of the prior precision signal occurs alongside the initial sensory mismatch, it concludes well before the extended processing tails of local adaptation and downstream model updates. This early resolution aligns directly with the precision’s prior-dependent role in weighting prediction errors - naturally subsiding alongside the surprisal response just as downstream model updating is initiated. Ultimately, these results demonstrate that these distinct model-derived signatures of adaptation and prediction differ in their latency profiles, temporally separating early mismatches from later model updates.

### 2.4 Repetition Suppression, Surprisal, Prior Precision and Model Updating Occupy Distinct Spectral Bands

Building on the temporal order of these predictive and adaptive effects, we next performed a post-hoc decomposition of the neural signals to investigate the specific spectral bands encoding these computations. To this end, the preprocessed MEG signals (at 250 Hz sampling) of each block were decomposed into seven distinct frequency bands: Delta (1–4 Hz), Theta (4–8 Hz), Alpha (8–13 Hz), Low Beta (13–20 Hz), High Beta (20–30 Hz), Low Gamma (40–70 Hz), and High Gamma (70–100 Hz). The lower frequency bands (Alpha to Beta) were computed using Morlet wavelets, while the higher frequency bands (Gamma bands) were estimated using a multitaper convolution approach. We then reran the identical time-shifted finite impulse response (FIR) regression independently on these band-filtered signals.

Spectral analysis was carried out in the same spatial clusters identified by the spatio-temporal broadband regression, isolating the specific spectral bands responsible for these effects. To avoid the mathematical limitations associated with projecting unsigned power through linear LCMV beamformers, this analysis was conducted in sensor space (see Methods 4.10). Specifically, we determined which frequencies drove the broadband effects by projecting their source-level statistical clusters through each participant’s forward model, selecting the top 15 most sensitive MEG sensors for each corresponding computational component. The signal-to-noise ratio within this spatial region was optimised by applying a singular value decomposition (SVD) across the 15 sensors and extracting the first principal component, upon which the band-specific Temporal Response Functions (TRFs) were subsequently analysed. To identify significant deviations in TRF power, we first employed a two-dimensional spectro-temporal cluster-based permutation test (spanning time and frequency). Multiple-comparison correction was applied to these 2D clusters using the False Discovery Rate (FDR) across the four components of interest, ensuring a highly conservative threshold for evaluating these spectral profiles. Additionally, because the sparse resolution of our spectral domain (limited to only seven discrete bands) might hinder the formation of robust spectro-temporal clusters, we subsequently performed one-dimensional temporal clustering independently within each spectral band. As this 1D approach corrects for multiple comparisons across time but not across the spectral dimension, significant findings from this secondary analysis are strictly denoted as statistical trends.

This spectro-temporal analysis revealed distinct frequency profiles for local adaptation versus statistical predictions, partially diverging from canonical spectral mappings. Following spectral decomposition, the repetition suppression effect was primarily observed as a reliable spectro-temporal cluster spanning the Theta and Alpha bands from 0 to 176 ms (*p*_FDR_ = .007), reflecting a dampening of spectral power. Additionally, a one-dimensional trend - corrected only across time within that specific band - indicated a further gain of power in the High Beta band from 128 to 224 ms (*p* = .026). While the theta modulation aligns with previously reported complex adaptation effects [27], the absence of gamma-band attenuation contrasts with classical models of repetition suppression [7]. In contrast, the spectral profile of surprisal was anchored by a reliable spectro-temporal cluster confined to the Low Beta band, extending continuously from 144 to 432 ms, characterised by a dampening of spectral power (*p*_FDR_ = .013). This dampening of (low-)beta power is consistent with the literature linking beta suppression to the reset of current internal models [26], whereas we did not observe a reliable increase within higher-frequency bands.

Higher-order predictive processes engaged frequencies classically associated with hierarchical routing [21]. For model updating, no reliable spectro-temporal clusters survived family-wise correction. However, a one-dimensional trend revealed an early transient dampening in High Gamma power from 64 to 176 ms (*p* = .020). As shown in the spectro-temporal data (Figure 4), this attenuation precedes the primary temporal peak observed in the broadband analysis. While gamma enhancements typically mediate feedforward model update signalling [22], the expected subsequent power gain corresponding to the main broadband response was absent. Consequently, this early dip may reflect a preparatory suppression prior to the primary model updating effect but must be interpreted cautiously given the lack of a sustained gamma increase and its restricted one-dimensional temporal correction. Finally, prior precision was characterised by a broad early spectral footprint, yielding two reliable spectro-temporal clusters. The first encompassed the Theta and Alpha bands from -48 to 224 ms (*p*_FDR_ = .022), while the second occurred concurrently in the High Beta band from -48 to 288 ms (*p*_FDR_ = .030); both clusters represented a dampening of spectral power. An additional one-dimensional trend suggested later modulation in the Low Beta band from 320 to 464 ms (*p* = .040). This widespread, multi-band oscillatory recruitment - particularly within the alpha and beta bands classically linked to top-down predictive state maintenance [31, 32] – aligns with prior precision’s role as a diffuse, modulatory mechanism scaling the integration of predictions and sensory evidence across distributed cortical networks.

**Figure 4:**
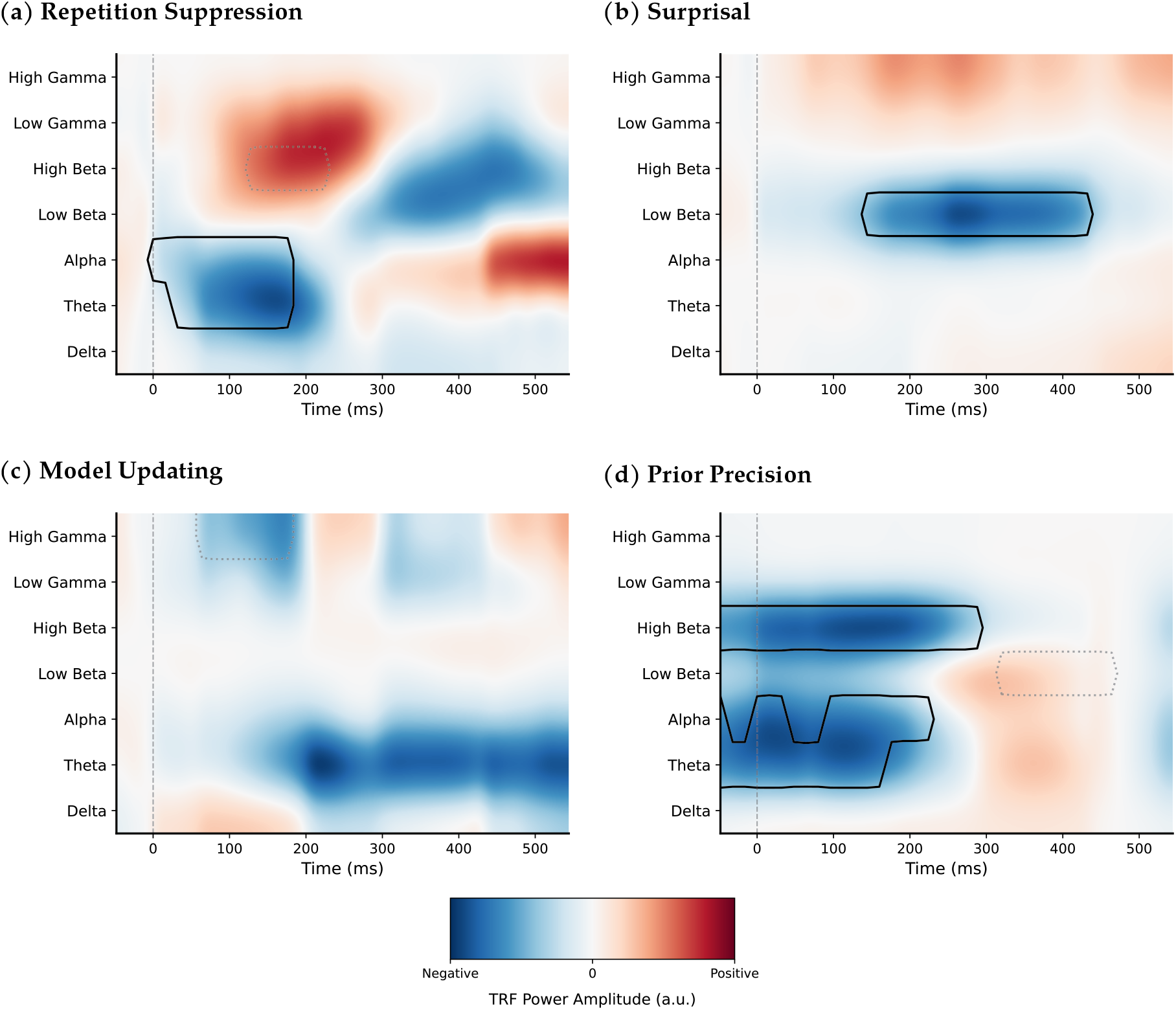
Distinct spectro-temporal signatures separate rapid sensory adaptation from hierarchical predictive processing. (a) Repetition suppression, (b) surprisal, (c) model updating, and (d) prior precision. Heatmaps display the band-specific temporal response functions (TRFs) extracted from source-guided MEG sensor arrays (top 15 most sensitive channels per component) and isolated via singular value decomposition (SVD). The y-axis represents the canonical frequency bands (Delta, Theta, Alpha, Low Beta, High Beta, Low Gamma, and High Gamma), and the x-axis indicates time relative to tone onset. Colour hue represents the cross-subject mean TRF power amplitude (arbitrary units). Amplitudes are scaled using an AUC-based weighting method to flatten spectral 1*/f* amplitude biases across frequency bands. Solid black contour lines delineate regions of statistical significance (*p*_FDR_ *<* 0.05), determined via non-parametric cluster-based permutation testing with FDR-correction. Dotted contour lines denote one-dimensional statistical trends - corrected only across time - within each specific frequency band. Heatmap arrays are visually smoothed utilising bicubic interpolation.

## 3 Discussion

By combining computational modelling with time-shifted FIR regressions of human MEG data, we investigated the temporal, spatial, and spectral dynamics of predictive and adaptive mechanisms within a continuously evolving auditory environment. We quantified the concurrent contributions of repetition suppression, surprisal, model updating, and prior precision. By isolating repetition suppression as an independent component acting outside the explicit hierarchical updates of predictive coding [11], we successfully disentangled local sensory adaptation from more involved statistical predictions. Our approach revealed a distinct chronological order: prior precision, driven by pre-stimulus expectations, was the earliest active component. This preceded the rapid (but lasting) low-level adaptation of repetition suppression, and the similarly timed, but more transient, mismatch detection of surprisal. These global mismatch evaluations were ultimately followed by delayed model updates. Spectrally, these components were likewise differentiable: repetition suppression was characterised by theta and alpha dampening, surprisal was confined to low-beta suppression, prior precision exhibited widespread multi-band dampening across theta, alpha, and beta bands, and model updating showed a transient trend toward high-gamma attenuation.

Crucially, delineating these dynamics highlights a functional and temporal divergence between early, stimulus-driven attenuation and downstream hierarchical model updating. While local repetition suppression and early surprisal operate on a rapid timescale within primary auditory cortices, they serve distinct roles: local adaptation attenuates sensory gain to recent acoustic history, whereas early surprisal acts as a global mismatch mechanism [19]. Spatially anchored in primary and associative sensory regions, this signal serves as a non-specific alert that prevailing expectations have been violated. In the context of dual-opponent accounts of perception [20], surprisal interrupts the brain’s initial bias toward expected structure, priming downstream networks to evaluate the exact nature of these deviations - prompting an ‘exploration’ state [33, 34]. As this signal transitions into later, distributed processing within frontoparietal networks, the brain shifts to active model revision. Together, this sequence illustrates how the brain coordinates fast, local sensory gating with slower, hierarchical error integration to continuously navigate stochastically evolving environments.

Running parallel to the initial detection of surprisal, the neural dynamics of repetition suppression largely mirror surprisal’s early temporal onset yet encompass wider cortical areas and exhibit a more sustained activation profile. Departing from surprisal, significant neural modulations associated with repetition suppression were evident even prior to stimulus onset and persisted extensively throughout the entire epoch. Such pre-stimulus presence reflects a dampening mechanism dependent on past acoustic history, while its broad distribution across bilateral temporal regions is characteristic of feedforward-driven sensory adaptation [35, 36]. Unlike explicit predictive updates, which require the continuous evaluation of inferential models, repetition suppression operates primarily as a local, parsimonious adjustment to the immediate sensory environment. This uniform profile matches recent mesoscopic evidence - employing an identical stochastic environment - showing that repetition suppression acts as a laminar-independent local gain control mechanism [11]. Consequently, this attenuation emphasises that the auditory system does not exclusively rely on the complex extraction of statistical regularities to navigate its environment. Rather, it concurrently depends on simpler mechanisms to efficiently gate redundant sensory information, thereby minimising computational load and aiding higher-order predictive processes.

Following these rapid sensory signatures, model updates peaked significantly later, near 190 ms, representing the context-informative shift from prior expectations to updated posterior states [12, 37]. To execute this revision, these delayed signals recruit a broader, distributed fronto-parietal network. Complementing this update process is precision, which functions as a continuous, ever-evolving metric closely linked to our current prior and is present even before stimulation onset times. This sustained, prior-dependent state mirrors the content-specific predictive models maintained in deep cortical layers during identical auditory environments [11]. However, its signal decay (178.7 ms) coincides with the resolution of the surprisal response (192.4 ms) and the onset of the model updating response (188.8 ms). This temporal alignment supports its theorised role in scaling prediction errors [17]: precision weighting resolves once sensory evidence has been integrated and subsides as model revision is initiated. Furthermore, prior precision displayed a unique spatial topology, engaging a widespread fronto-temporal network alongside a distinct, late-emerging cerebellar cluster. This cerebellar involvement aligns with its established role in forward modelling and the continuous fine-tuning of internal predictive states [38, 39, 40]. Given that prior precision effects are maintained during the earlier timeframe of surprisal, it remains an open question whether this global mismatch detection is similarly scaled by precision.

Furthermore, analysing the data separately across each frequency band allowed us to isolate the specific spectro-temporal dynamics of these components, which partially diverged from canonical spectral mappings. For repetition suppression, we observed significant theta and alpha dampening alongside a trend in the high-beta band, but lacked the characteristic gamma-band attenuation often ascribed to rapid sensory adaptation [7]. We propose two distinct reasons for this absence. First, splitting the data into its constituent frequency bands severely reduced the signal-to-noise ratio (SNR) of the Temporal Response Functions (TRFs), compounding the inherent difficulty of capturing high-frequency gamma signals reliably in MEG. Time-frequency representations did indicate a cluster starting at high beta that might encompass low gamma, but our current analysis lacked the resolving power to extract it statistically. Second, because we modelled repetition suppression concurrently with hierarchical computations, part of the high-gamma variance normally ascribed to repetition suppression in isolation may now be uniquely captured by the model updating parameter. This SNR limitation likely extends to our estimates of surprisal; while we observed the expected dampening in the beta band [26], a visually apparent positive divergence in the high-gamma band failed to reach significance. Similarly, for model updating, while we detected an early statistical trend in high-gamma dampening, stringently controlled spectral results - such as the gamma enhancements classically linked to model updating [22] — remained absent. Finally, prior precision exhibited a broad spectral footprint, dampening theta, alpha, and beta bands, aligning with its modulatory role in maintaining predictive states. While our broadband framework successfully separated these parameters in the time domain, a paradigm explicitly optimised for time-frequency analysis and higher SNR is warranted to better disentangle the underlying spectral components of these mechanisms while still modelling them concurrently and dynamically.

Ultimately, these findings indicate the methodological necessity of evaluating adaptive and predictive mechanisms concurrently. Because top-down statistical predictions frequently coincide with the repetition of low-level acoustic features, paradigms that investigate mechanisms like repetition suppression or mismatch negativity in isolation risk conflating localised sensory adaptation with higher-level predictive updating [7, 11]. Although these mechanisms share overlapping features - such as the early latencies and auditory recruitment of repetition suppression and surprisal - they ultimately exhibit partially dissociable spatial, temporal, and spectral profiles. This suggests that previously reported unitary effects likely represent a blended integration of distinct hierarchical processes. These temporal dissociations complement recent laminar fMRI evidence from the same paradigm, showing that while predictive models and prediction-input alignments are confined to deep and superficial layers respectively, repetition suppression acts as a depth-invariant gain control. It must be noted, however, that the exact spatial boundaries of these overlapping networks remain constrained by spatial leakage inherent to MEG beamforming. Furthermore, because our computational framework relies on specific model hyperparameters and a passive listening paradigm, future validation is required to determine how precisely these isolated neural signatures translate to active perception.

Auditory perception is not a singular operation, but a cascade of different processes. It relies on the brain continuously balancing immediate, local acoustic adaptation with the complex integration of overarching statistical contexts. By mapping this sequential progression - from rapid, uninformative sensory mismatch detection to distributed, precision-weighted model updates - our findings provide a framework for how the brain navigates and parses a continuously evolving acoustic world.

## 4 Methods

### 4.1 Participants

Thirty-three participants (eleven male;*M*= 25.5 years, age range: 19–44 years) participated in this magnetoencephalography (MEG) study. All participants reported normal hearing and no history of neurological disorder. The study was approved under the general ethical approval for the Donders Centre for Cognitive Neuroimaging (Imaging Human Cognition, CMO2014/288) by the local ethics committee (CMO Arnhem-Nijmegen, Radboud University Medical Centre). Participants provided written informed consent and received monetary compensation for their participation. One participant was excluded due to data corruption. The final sample, therefore, consisted of 32 participants.

### 4.2 Experimental Setup

Participants were seated in a dimly lit, magnetically shielded MEG room (MSR) and wore a neck brace for head stabilisation. To minimise movement artefacts, they were instructed to maintain a stable head position and fixate on a central crosshair (85 cm viewing distance) while passively listening to the presented tones. Head position was continuously monitored using fiducial markers on the nasion and the left and right preauricular points [41]. Between blocks, participants readjusted their head position if it deviated from the starting point by more than 5 mm. Gaze was concurrently tracked using an SR Research Eyelink 1000.

Stimuli were presented using MATLAB 2023b (MathWorks Inc., Natick, USA) and the Psychophysics Toolbox [42, 43, 44]. Mono-audio was presented over loudspeakers in the MSR. Prior to the main experimental runs, participants completed a calibration task to ensure consistent perceived loudness across the frequency spectrum. During this task, participants compared pairs of tones across nine calibration frequencies, logarithmically spaced between 200 Hz and 6,000 Hz. These comparisons were subsequently used to generate individualised equal-loudness curves [45]. Intermediate values were estimated using piecewise cubic interpolation to create a continuous curve, which was then applied to adjust the intensity of the experimental tones.

Participants were permitted breaks between blocks and used an MEG-compatible button box to resume testing.

### 4.3 MEG and MRI Acquisition

MEG data were acquired using a 275-channel axial gradiometer CTF-MEG system (CTF MEG Systems, Coquitlam, Canada). The continuous signals were digitised at a sampling rate of 1200 Hz, and low-pass filtered online at 300 Hz. Between zero and three channels were disabled prior to recording, leaving 272–275 active channels per participant.

The participant’s head shape and the locations of three fiducial points were digitised using a Polhemus 3D tracking device. The MEG data were complemented by T1-weighted anatomical MRI scans acquired on a 3T Siemens system using an MPRAGE sequence (TR = 2.3 s, TE = 3.03 ms, flip angle = 8°, GRAPPA acceleration factor = 2, 192 transversal slices, 1 mm isotropic voxel size).

### 4.4 Experimental Design

Auditory sequences comprised pure tones sampled probabilistically from a mixture of two Gaussian distributions (defined on alog _2_-frequency axis). Tones were drawn from one of four distinct Gaussian pairs with centre frequencies ranging from 500 Hz to 3,000 Hz: (500/1392 Hz), (646/1790 Hz), (834/2323 Hz), and (1,078/3,000 Hz). To ensure perceptual distinguishability, Gaussian centres within each pair were separated by 1.48 octaves. Each distribution had a fixed bandwidth of 1/3 octave (FWHM), with tones sampled at a 1/12th octave resolution to yield uniformly spaced candidate frequencies.

The approximately one-hour experimental session consisted of 12 runs, each subdivided into two blocks containing 10 stable periods. Within each stable period, tones were sampled with fixed probabilities from the two underlying distributions (A and B), preserving local statistical regularity. The relative sampling probability between the two distributions was manipulated across periods using a logistic transformation, yielding 15 distinct steps ranging from 5% to 95%. These probabilities were assigned pseudo-randomly to ensure broad coverage while avoiding immediate repetitions of extreme values. Consequently, transitions between stable periods introduced statistical shifts of varying magnitude, ranging from perceptually salient (e.g., 95% A to 95% B) to subtle (e.g., 95% A to 90% A). Because sampling was probabilistic, the perceptual clarity of these transitions varied throughout the runs.

Tones were presented for 200 ms with a 50 ms inter-tone interval (250 ms stimulus onset asynchrony). Each stable period featured 48 tones (12 seconds) before the sampling probability changed, resulting in 11,520 tones per participant. Finally, due to severe MEG artefacts, some blocks were excluded from analysis: four participants retained 23 of 24 blocks, two retained 22, and one retained 18.

### 4.5 Computational Modelling

Computational modelling was applied on each tone in the stochastic sequences to estimate how repetition suppression, prior precision, surprisal, and model updating evolve over time. Normative estimations were obtained using the computational models described below.

#### 4.5.1 Long-trace Repetition Suppression Model

We employed a double-exponential model of long-trace repetition suppression, motivated by evidence that brief sensory stimulation induces persistent, stimulus-specific suppression within early sensory cortices [8]. This model posits that the residual effect of a stimulus presented *n* trials prior decays over time according to two exponential components, capturing distinct short- and long-lasting suppressive mechanisms. Formally, the repetition suppression factor *f* (*n*) is expressed as:

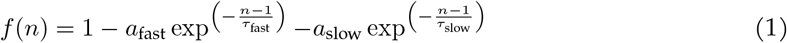

where *a*_fast_ and *a*_slow_ define the amplitudes of the fast and slow suppressive components, and *τ*_fast_ and *τ*_slow_ represent their corresponding time constants, expressed in trials. Parameter values were adopted from a previous neurophysiological investigation in primary visual cortex (V1) [8] and were recently shown to be consistent with fMRI responses in the auditory cortex using the exact same stimulus paradigm [11]. Specifically, we set *a*_fast_ = 13.99% with *τ*_fast_ = 0.85 trials, and *a*_slow_ = 3.45% with *τ*_slow_ = 6.82 trials. These parameters describe two temporally separable phenomena: an acute phase of rapid adaptation and a more prolonged, sustained suppressive state.

To estimate cumulative suppression across these dual timescales, the effective response of each frequency unit to an incoming stimulus was scaled by the combined influence of the preceding ten trials. Because this suppressive effect is stimulus-specific, the *n*-back decay vector weights the temporal history of prior tones independently for each frequency channel. By projecting the sequence of presented tones through this decay profile, we established a forward model that dynamically tracks parallel, frequency-specific adaptation states. Rather than relying on a global adaptation metric, this approach inherently accounts for continuous frequency selectivity, ensuring that adaptation to a specific tone persists accurately within its tuned population.

#### 4.5.2 D-REX Model

The Dynamic Regularity Extraction (D-REX) model employs Bayesian sequential prediction with perceptual constraints, primarily focusing on memory (*m*), to simulate the brain’s processing of sound sequences over time [28]. Memory (*m*) restricts the number of prior time points considered; here, we set *m* to be infinite within the boundaries of a block, ensuring the model integrates the continuous context of that entire block uninterrupted by pauses. The model continuously predicts the distribution of the next time point (*x*_*t*+1_) based on previous observations (*x*_1:*t*_) by estimating local statistics (*θ*), corresponding to the sample mean and variance of the input.

In addition to these features, the model incorporates a Gaussian Mixture Model (GMM) to represent multiple components within the sound sequences. For our stimuli specifically, we constrained the maximum number of components to two, corresponding to the two centre frequencies present in the sequence of stimuli. To manage the formation of new components within the GMM, a threshold parameter,*β*, was set to 0.2. This*β*value, scaled by predictive probabilities, further constrains the overall stimulus space.

The model accounts for possible changes in parameters(*θ*)by generating hypotheses across varying run lengths(*r* _*t*_)and integrating these to predict future observations:

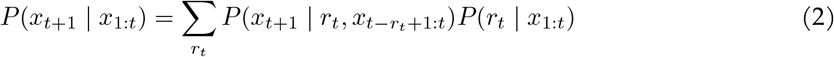

The primary output of the model is surprisal(*S* _*t*+1_), representing the discrepancy between the predicted probability and the actual observation:

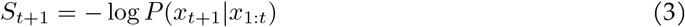

Lower probability events yield higher surprisal, while zero surprisal corresponds to perfectly predicted events. Additionally, the model computes prior precision, defined as the inverse of the variance across the full predictive distribution, thereby quantifying the confidence in the prior predictions over time:

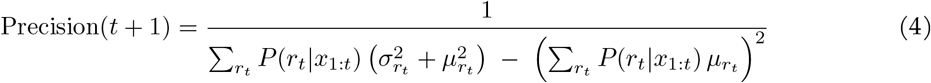

Here, *P*(*r* _*t*_ | *x* _1:*t*_)is the posterior probability of a run length *r*_*t*_ given observations up to time *t*, while 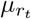 an 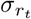 denote the predicted mean and standard deviation, respectively, of the next stimulus under run length *r* _*t*_.

#### 4.5.3 Model Updating

Beyond computing surprisal and prior precision for each observed stimulus, the D-REX model estimates the prior probability of each frequency across the entire stimulus space. This continuous probability distribution explicitly reflects the model’s expectations based on previously encountered auditory regularities. To quantify model updating at each time point using these inferred predictive distributions, we adopted an area-based metric conceptually aligned with recent predictive coding frameworks [29]. We defined model updating as the degree of non-overlap between the normalised probability distributions of the current posterior and the prior expectations. Formally, the model updating (*MU*) at time step *t* was calculated as half the integrated absolute difference between the two distributions:

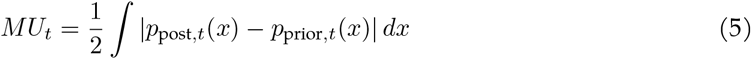

Where *x* represents the relevant feature space (i.e., frequency domain). The scaling factor of0.5 normalises the *MU* to a bounded range of [0, 1], where0indicates identical, perfectly overlapping distributions (zero model update) and1indicates completely non-overlapping distributions (maximal model update).

### 4.6 MEG Analysis

#### 4.6.1 Pre-processing

All MEG data were pre-processed in MATLAB 2023b using FieldTrip (2023 release; [46]). Raw data were loaded per run, defining trials with 2.0 seconds of pre-run and 3.0 seconds of post-run periods based on stimulus triggers. Relying on the reference sensors of the CTF MEG system, we denoised the recorded MEG data using third-order gradient correction, followed by demeaning. The continuous run data were then segmented into one-second chunks. Using visual artefact rejection routines in FieldTrip (i.e.`’summary’`), we removed noisy segments and sensors exhibiting excessive variance (e.g., SQUID jumps). We repeated this rejection procedure for muscle artefacts using data bandpass-filtered between 110 and 140 Hz (4th-order Butterworth filter). After removal of noisy segments, the data were resampled to 400 Hz. Independent component analysis (ICA, utilising `‘runica’`) was performed on the combined data for all runs per participant to identify components reflecting artefacts such as cardiac activity, residual eye movements, or blinks. Following ICA, the continuous unsegmented data were reprocessed (gradient correction and demeaning), and bad components were removed. MEG timestamps were explicitly synchronised using the system’s clock channel (SCLK01). Finally, the clean data and trigger channels were downsampled to 60 Hz for subsequent analysis. This clean data was also downsampled in the same way to 250 Hz for the spectro-temporal analysis.

#### 4.6.2 Finite Impulse Response (FIR) Model

We applied time-resolved linear regression to the MEG data to explore the temporal dynamics of tonal repetition suppression, surprisal, model updating, and prior precision, employing a time-shifted finite impulse response (FIR) model (rERP; [47]). This approach explicitly disentangles responses to concurrent tonal features and corrects for continuous temporal overlap from adjacent stimuli. Each feature of interest was assigned a single continuous value per tone (*x ∈* ℝ), which was subsequently expanded into a time-shifted FIR design matrix to estimate the temporal response function at the 60 Hz sampling resolution of the MEG data. To optimise this deconvolution and capture non-stimulus variance, the design matrix incorporated nuisance regressors accounting for the inter-stimulus interval (ISI), block status, adjacent tone onsets and offsets, a global constant, and 3rd-order polynomial drifts per block. MEG data were z-scored separately for each run, and regressors were normalised to range from 0 to 1.

We constructed a time-expanded design matrix **M**, including time-shifted versions of each regressor arranged column-wise. The temporal window spanned from*t* _*min*_ = *−* 50ms to*t* _*max*_ = 567 ms relative to tone onset. This specific upper bound was selected so that the window length in samples (at a 60 Hz sampling rate) is coprime with the 250 ms stimulus presentation period, ensuring an optimal balance of distances to adjacent tone onsets and offsets and minimising periodic stimulus biases. This yielded 37 delayed columns per regressor.

Next, we estimated the regression weights 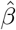 using ordinary least squares regression:

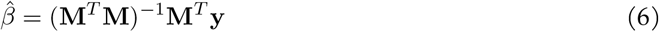

Throughout this estimation, we employed a leave-one-block-out cross-validation procedure, using the mean beta coefficients across folds for subsequent analysis.

Collectively, the beta weights constitute a temporal response function (TRF), depicting how features modulate neural activity over time [48, 49]. In practice, the TRF reflects how parametric variations in a predictor are expressed in MEG-sensitive neural activity at different latencies.

To generate a null distribution, we recalculated the same FIR - OLS - regressions using a permuted design matrix (**M**). The exact temporal alignment between the neural data and all nuisance regressors was maintained to properly account for baseline physiological noise. For the target TRF regressors, we applied a random block-wise permutation (swapping intact blocks of predictors) followed by a random circular shift (e.g., 15–25 s) within the block. This strategy preserves the temporal autocorrelation and covariance structure of the predictors while completely decoupling their temporal phase from the neural data, establishing a strict and unstructured noise baseline.

### 4.7 Source Estimation

Individual T1-weighted anatomical MRIs were co-registered to the MEG coordinate system using three fiducial points and further refined by fitting the digitised Polhemus headshape to the MRI-extracted scalp surface. For one participant lacking an anatomical MRI, a standard template MRI was utilised and affinely warped to match their individual digitised Polhemus headshape. A single-shell volume conduction model [50] was then created for each participant. Subject-specific source grids with a 0.5 cm resolution were generated by non-linearly warping a standard MNI template grid to the individual anatomies [46]. Leadfields were computed for each grid point, incorporating the third-order gradient balancing of the CTF system and applying a rank reduction of two to account for the physical properties of the MEG signal.

Source reconstruction was performed utilising a Linearly Constrained Minimum Variance (LCMV) beamformer [51]. To enable direct comparisons across the different computational predictors, a common spatial filter was computed for each participant. The data covariance matrix used to construct this filter was derived from the continuous, pre-processed sensor data, which was segmented into 3-second overlapping pseudo-epochs to ensure stable covariance estimates. Data rank was explicitly evaluated, and`’Tikhonov’`regularisation was applied, with the regularisation parameter selected individually for each participant based on the knee of the spectrum. To correct for the inherent depth bias of the beamformer, the filter weights were normalized utilizing a unit-noise gain, yielding a Neural Activity Index [52]. Finally, the sensor-level time-resolved FIR regression weights (TRFs) as well as their null counterparts were multiplied by this common spatial filter to project the temporal dynamics of each predictor into source space.

### 4.8 Polarity Alignment

In the source-localised MEG data, the reconstructed dipole moments in individuals are symmetric in polarity, with the different sources having an arbitrary sign due to the inherent orientation ambiguity of the SVD-based common filters. To harmonise the polarities across participants, we performed a data-driven polarity alignment procedure.

This harmonisation was achieved by applying an iterative singular value decomposition (SVD) to the source-level timecourses. Specifically, the sign of each source was flipped in accordance with its polarity relative to the first singular vector. Because this vector represents the dominant spatial correlation across the cortex, this step effectively aligns the phase of all regional responses to the dominant functional network. To resolve the remaining global sign ambiguity and ensure uniformity across the cohort, the entire dataset was anchored to its strongest response; the harmonised matrix was uniformly multiplied by the sign of the absolute maximum peak, forcing the primary temporal deflection to be strictly positive.

Crucially, while this procedure enforces cohort uniformity, the eventual global sign remains biologically arbitrary. Consequently, this alignment was utilised to prevent the phase-cancellation of group-average waveforms during visualisation. All primary statistical evaluations, including the spatiotemporal clustering, were performed on the squared power of the source signals, ensuring the core formal analyses remained entirely independent of dipole polarity.

### 4.9 Spatio-temporal Clustering

To evaluate the statistical significance of the source-localised temporal response functions, we conducted non-parametric spatiotemporal permutation testing. Empirical TRF source power was compared against the unstructured null baseline utilising a one-tailed dependent-samples*t*-test across the full continuous modelled temporal window ( *−* 50ms to567ms). To effectively control for multiple comparisons across space and time without imposing arbitrary initial cluster-forming thresholds, we applied Threshold-Free Cluster Enhancement (TFCE; [53]) using standard spatial integration parameters (*H*= 2.0,*E*= 0.5). The empirical null distribution was approximated via 2,000 random permutations of the true and null condition labels. Finally, to rigorously control the family-wise error rate across the evaluation of the four distinct computational predictors (repetition suppression, surprisal, model updating, and prior precision), the ultimate cluster-level significance threshold was strictly bounded at*α*= 0.01.

#### 4.9.1 Functional Region of Interest Selection

To facilitate the visualisation of the temporal response functions (TRFs), we implemented a data-driven region of interest (ROI) selection procedure. First, to mitigate the influence of transient spatial noise, we applied a 100 ms rolling temporal window within the boundaries of each significant spatiotemporal cluster. Within each window, the 100 most active voxels received a vote. This iterative voting system isolated a core functional network of robustly activated voxels, which we then used to extract the overarching temporal peak of the cluster. To anchor the cluster anatomically, its full spatial extent was intersected with a standard anatomical atlas (AAL; [54]). By calculating the total statistical mass (area under the curve) of the cluster, we determined its proportional distribution across the underlying anatomical parcels; the parcel capturing the highest percentage of this mass was formally designated as the representative spatial ROI. Finally, to quantify the broader spatial extent of each network, we computed two additional metrics strictly within the same 100 ms window centred on the extracted temporal peak. The ‘peak footprint’ was defined as the percentage of the modelled intracranial grid surviving TFCE correction (*p <*0.01) during this peak timeframe. Concurrently, ‘anatomical dispersion’ was calculated as the minimum number of distinct AAL parcels required to capture 80% of the cluster’s integrated TFCE mass within this same window.

### 4.10 Spectral-TRF Disentanglement

To investigate the frequency-domain correlates of prediction and the complementary adaptive mechanism of tonal repetition suppression, we extended the identical cross-validated FIR modelling approach described above to extract spectro-temporal response functions (Spectro-TRFs) from distinct spectral bands. First, continuous neuromagnetic responses were segmented into block-wise arrays (i.e., isolated continuous task segments of 480 tones) and notch-filtered at 50 Hz and 100 Hz to attenuate line noise.

We decomposed these neuromagnetic responses into canonical frequency bands using two distinct time-frequency transformation strategies optimised for low and high frequencies, maintaining a 250 Hz sampling rate. For low-frequency bands-delta (1–4 Hz), theta (4–8 Hz), alpha (8–13 Hz), and beta (low: 13–20 Hz, high: 20–30 Hz), we computed spectral power using Morlet wavelets with the number of cycles defined dynamically as*f/*2. For high-frequency gamma bands (low: 40–70 Hz, high: 70–100 Hz), we utilised discrete prolate spheroidal sequences (DPSS) multitaper convolutions to optimise spectral concentration and manage high-frequency variance. Multitaper parameters included a frequency step resolution of 5 Hz, a time-bandwidth product of 5, and an adaptive window length proportional to 20% of the target frequency. Power estimates were subsequently averaged across frequency bins within each defined band.

Following spectral decomposition, blocks exhibiting excessive noise within any of the spectral components were disregarded (yielding an average of 20.56 retained blocks per participant,*SD*= 2.61). The remaining band-specific power fluctuations were modelled as dependent variables within the ordinary least squares (OLS) regression framework. To mitigate high-frequency temporal noise and maintain a temporal resolution like the broadband TRF analysis, the frequency data were analysed by binning the regressors into 16 ms temporal windows (compared to the 16.67 ms temporal bins within the broadband FIR regression). To align with the 250 Hz sample grid, the modelled temporal window was adjusted to span from *t* _*min*_ = *−* 48ms to*t* _*max*_ = 560ms relative to tone onset (compared to the *t* _*min*_ = *−* 50ms to*t* _*max*_ = 567ms of the broadband FIR regression). This configuration yielded 38 delayed response bins per regressor, ensuring the resulting Spectro-TRFs maintained an approximately equivalent parameter space and comparable temporal resolution to the primary broadband FIR model.

To avoid the mathematical limitations associated with projecting unsigned power through linear spatial beam-forming filters, all spectral analyses were conducted in sensor space, spatially guided by the preceding broadband findings. Specifically, we utilised the spatial topographies identified in the broadband spatio-temporal source analysis to isolate the MEG sensors that maximally captured activation within these clusters. For each participant, source-level statistical masks - derived from the statistical mass (AUC) of the significant TFCE clusters - were multiplied by the individual’s forward model (leadfield matrix). This procedure enabled us to rank the sensors based on their physical sensitivity to the underlying neural sources, allowing us to select the top 15 most sensitive sensors for all subsequent statistical analyses. To maximise the signal-to-noise ratio and derive a single representative time course per participant, we applied a singular value decomposition (SVD) across these sensor subsets prior to statistical testing.

### 4.11 Statistical Testing

To systematically evaluate temporal differences in component latencies among the computational predictors, we performed jackknife-based latency *t*-tests [30]. Component latencies were quantified using a 50% relative amplitude (half-maximum) criterion, restricted to the significant spatiotemporal windows established by the preceding cluster analysis. All *p*-values derived from these pairwise temporal comparisons were subsequently corrected for multiple comparisons using the False Discovery Rate (FDR) procedure.

To evaluate the statistical significance of the resulting Spectro-TRFs, we employed a non-parametric cluster-based permutation testing framework operating across the two-dimensional time-frequency space. For each computational regressor, individual participant data were defined by the 7 canonical frequency bands and 38 discrete temporal bins. A one-sample*t*-statistic against zero was computed at each band-by-time coordinate. Contiguous data points exceeding an uncorrected threshold were grouped into two-dimensional clusters, and a cluster-level test statistic was derived by calculating the mass (i.e., the sum of the *t*-values) within each bounded cluster. To control for family-wise error rate across the 2D parameter space, we constructed an empirical null distribution via a Monte Carlo randomisation procedure. Data signs were iteratively permuted across 10,000 iterations, extracting the maximum cluster statistic at each step. Final *p*-values for the observed Spectro-TRF clusters were defined as the proportion of randomised partitions that yielded a maximum cluster mass exceeding that of the original, unpermuted data. All resulting 2D cluster *p*-values were subsequently corrected for multiple comparisons over the four components of interest using hierarchical False Discovery Rate (FDR) [55]. Additionally, to account for instances where the independent dynamics of distinct frequency bands precluded the formation of reliable two-dimensional clusters, Spectral TRFs were also evaluated independently per band. This one-dimensional analysis mirrored the permutation procedure described above (10,000 iterations) but clustered contiguous data points exclusively across the 38 temporal bins within each of the 7 frequency bands. Because the resulting cluster *p*-values from this approach inherently correct for multiple comparisons across time but not across the distinct spectral dimension, significant 1D clusters are strictly denoted as statistical trends.

## 4.12 Acknowledgements

This work was supported by the European Research Council (ERC) under the European Union’s Horizon 2020 research and innovation programme (grant agreement No. 101001270).

## 4.13 Author Contributions

Conceptualisation: JJGvH. Data wrangling and preprocessing: JJGvH. Formal analysis: JJGvH. Statistical analysis and visualisation: JJGvH. Supervision: FdM, SAK, FPdL. Initial draft: JJGvH. Final draft: JJGvH, SAK, FPdL, FdM.

## 4.15 Data and Code Availability

All data, preprocessing code, modeling code and analysis scripts are freely available from the Donders Repository under CC-BY-4.0 license, under identifier di.dccn.DSC_3018063.01_731.

