## Supplementary materials for "Temporal Progression of Prior Precision, Surprisal, Model Updating and Tone Adaptation in Human Auditory Cortices"

### A Supplement

| **Predictor** | Surprisal | Model updating | Prior precision | VIF | Unique Var. |
|---|---|---|---|---|---|
| Repetition suppression | 0.68 | 0.29 | 0.25 | 1.93 | 52% |
| Surprisal | – | 0.41 | 0.16 | 2.05 | 49% |
| Model updating | – | – | 0.22 | 1.24 | 81% |
| Prior precision | – | – | – | 1.10 | 91% |

**Table S1**: **Correlations and Collinearity of Computational Predictors.** This table reports the absolute Pearson correlation coefficients ($|r|$) as well as the Variance Inflation Factor (VIF) between the four computational predictors included in the time-shifted FIR deconvolution. Values are derived from the stochastic stimulus sequences. The stochastic stimulus generation was explicitly designed to restrict collinearity (e.g., maintaining $r \approx 0.68$ between repetition suppression and surprisal, compared to near-perfect correlations in classical oddball paradigms). To confirm the full model successfully decorrelated these mechanisms, we also report the VIF and its inverse, Unique Variance (Tolerance). VIF quantifies how much a predictor overlaps with a combination of all other predictors combined. The Unique Variance reflects the percentage of a component's information that is completely orthogonal to the rest of the model. All predictors yielded VIFs well below the canonical threshold of 5 (max VIF = 2.05), demonstrating that at least 49% of the variance for every parameter was unique. This ensures the deconvolution framework could independently resolve these distinct neural signatures without misattribution.

| **Predictor** | Onset (ms) | Peak (ms) | Offset (ms) |
|---|---|---|---|
| Prior precision | -12.8 | 167.0 | 178.7 |
| Surprisal | 80.2 | 117.0 | 192.4 |
| Repetition suppression | 74.1 | 183.0 | 311.0 |
| Model updating | 188.8 | 200.0 | 398.3 |

**Table S2**: **Temporal Dynamics of Predictive and Adaptive Computations.** This table reports the temporal progression of the modelled neural responses. Onset and offset latencies reflect the local fractional latencies (50% relative amplitude criterion) derived from the jackknifed estimates, marking the transient emergence and resolution of each computational process. Peak latencies correspond to the maximum statistical strength of the significant TFCE clusters. Prior precision represents a pre-existing top-down state, reaching its relative half-maximum prior to stimulus onset (-12.8 ms).

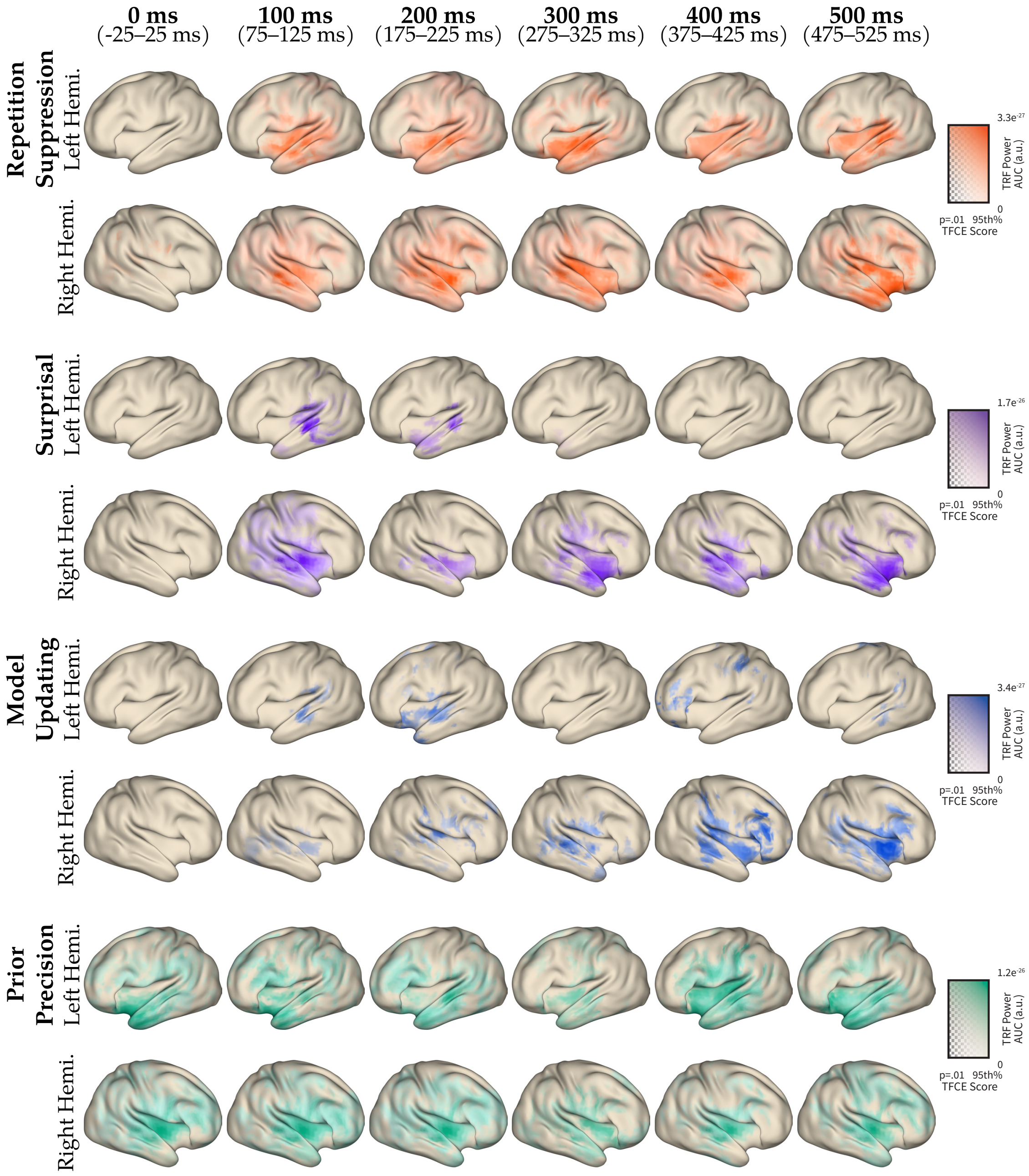


**Figure S1**: **Spatiotemporal evolution of cortical power responses.** Cortical surface maps display the distribution of source-localised MEG power for four experimental components: repetition suppression, surprisal, model updating, and prior precision. Rows alternate between Left and Right Hemisphere views for each component. Columns represent snapshot time windows centred at 0, 100, 200, 300, 400, and 500 ms post-stimulus. Each snapshot averages data over a 50 ms integration window (exact ranges indicated below column headers). The displayed metric is the area under the curve (AUC) of the power response within the respective time window. Colorbar scales indicate the magnitude ranges specific to each component's statistical map, matched across all time points within that condition.

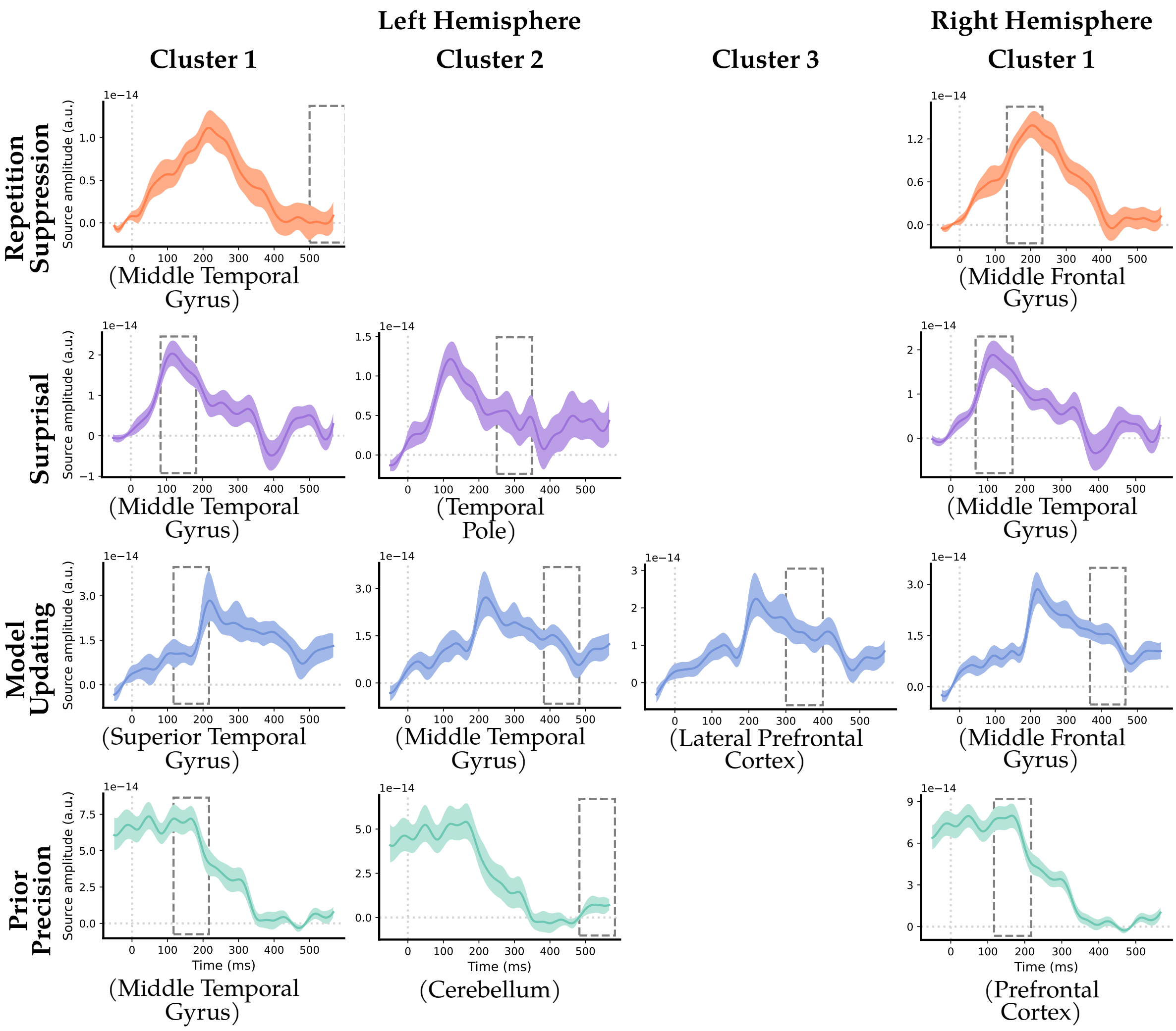


**Figure S2**: **Stability of temporal response functions across independent hemispheric analyses.** Polarity-aligned temporal response functions (TRFs) are shown for all significant spatiotemporal clusters derived when performing the Threshold-Free Cluster Enhancement (TFCE) permutations separately for the left and right hemispheres. Investigating all significant constituent clusters independently yielded similar waveform dynamics, demonstrating the stability and robustness of the results. Columns represent the individual significant clusters (up to three for the left hemisphere and one for the right hemisphere) for repetition suppression, surprisal, model updating, and prior precision.

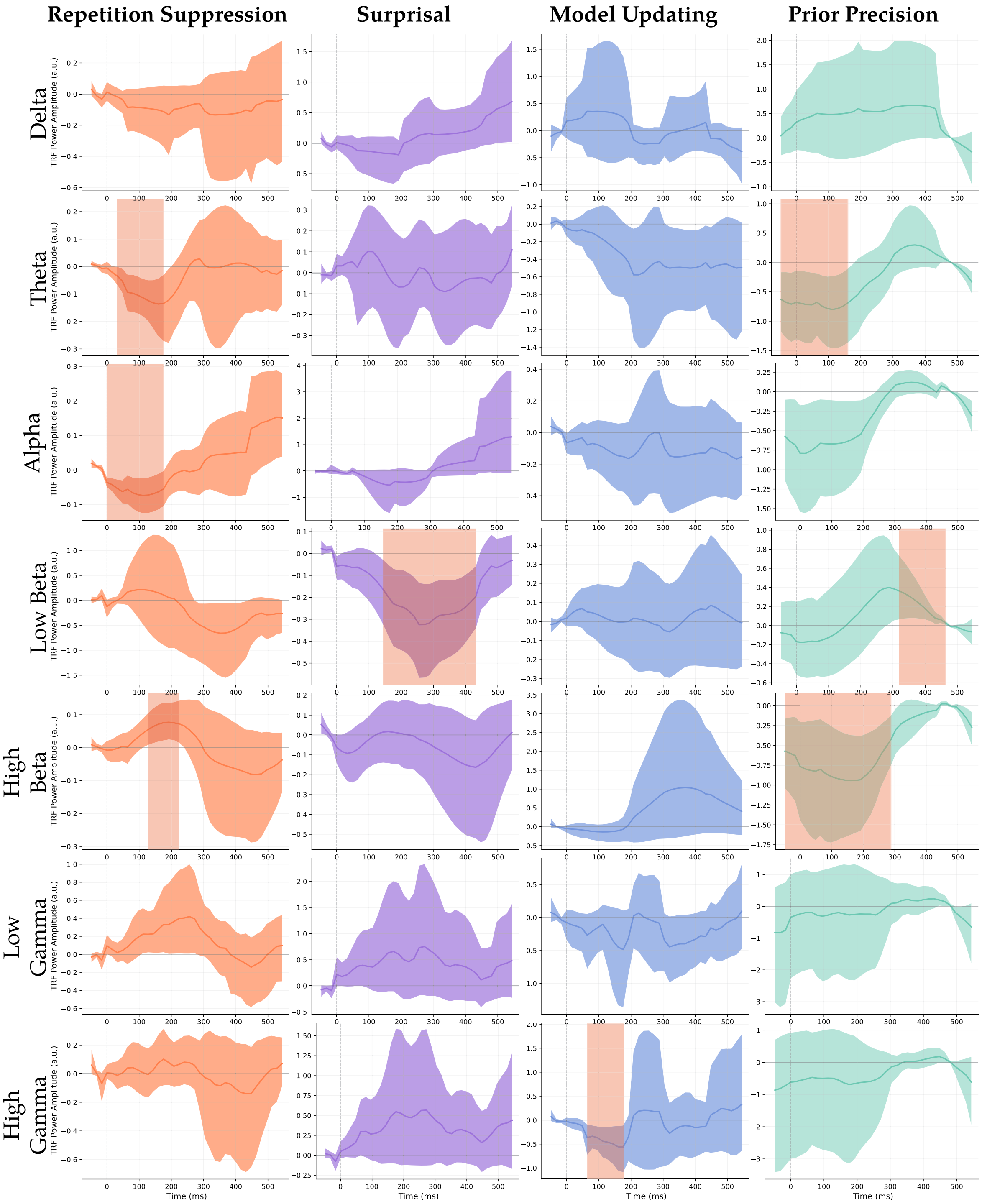


**Figure S3**: **Band-specific temporal response functions across computational components of sensory adaptation and predictive processing.** (a) Repetition suppression, (b) surprisal, (c) model updating, and (d) prior precision. Grids display the isolated temporal response functions (TRFs) extracted from source-guided MEG sensor arrays (top 15 most sensitive channels per component) via singular value decomposition (SVD) for each canonical frequency band. Rows represent canonical frequency bands (Delta, Theta, Alpha, Low Beta, High Beta, Low Gamma, and High Gamma), and columns indicate regression components. The x-axis indicates time relative to tone onset, and the y-axis represents cross-subject mean TRF power amplitude (arbitrary units), with shaded regions denoting 95% bootstrap confidence intervals. Amplitudes are scaled using an AUC-based weighting method to flatten spectral $1/f$ amplitude biases across frequency bands. Red shaded areas represent regions of statistical significance ($p < 0.05$), determined via non-parametric cluster-based permutation testing against zero.

| **Predictor** | Peak Window (ms) | Footprint (% grid) | Dispersion (AAL parcels) |
|---|---|---|---|
| Prior precision | 117–217 | 91.6 | 57 |
| Surprisal | 67–167 | 31.0 | 22 |
| Repetition suppression | 133–233 | 77.8 | 48 |
| Model updating | 150–250 | 37.7 | 45 |

**Table S3**: **Spatial Footprints and Anatomical Extent of Predictive and Adaptive Computations.** This table reports the spatial extent of the modelled neural responses. Spatial metrics were assessed strictly within a 100-ms window centred on each component's maximum peak. Prior precision displayed the most dispersed profile, followed by repetition suppression. While surprisal and model updating exhibited similarly restricted overall spatial footprints, their regional distributions differed: surprisal was more concentrated, whereas model updating extended across a broader array of AAL parcels, indicative of a more distributed network. Total footprint indicates the percentage of the modelled intracranial grid surviving TFCE correction ($p < 0.01$). Anatomical dispersion reflects the number of distinct AAL parcels required to capture 80% of the integrated TFCE mass within that footprint. Due to spatial leakage inherent to MEG beamforming, footprint percentages serve only as a relative scale to compare spatial recruitment between conditions rather than absolute boundaries of neural tissues.
